# Using transcriptome sequencing and pooled exome capture to study local adaptation in the giga-genome of *Pinus cembra*

**DOI:** 10.1101/462630

**Authors:** Christian Rellstab, Benjamin Dauphin, Stefan Zoller, Sabine Brodbeck, Felix Gugerli

**Affiliations:** WSL Swiss Federal Research Institute, Zürcherstrasse 111, 8903 Birmensdorf, Switzerland; ETH Zürich, Genetic Diversity Centre, Universitätstrasse 16, 8092 Zürich, Switzerland

**Keywords:** Conifer, exome capture, paralogs, pine, Pool-Seq, transcriptome assembly

## Abstract

Despite decreasing sequencing costs, whole-genome sequencing for population-based genome scans for selection is still prohibitively expensive for organisms with large genomes. Moreover, the repetitive nature of large genomes often represents a challenge in bioinformatic and downstream analyses. Here we use in-depth transcriptome sequencing to design probes for exome capture in Swiss stone pine (*Pinus cembra*), a conifer with an estimated genome size of 29.3 Gbp and no reference genome available. We successfully applied around 55,000 self-designed probes, targeting 25,000 contigs, to DNA pools of seven populations from the Swiss Alps and identified > 140,000 SNPs in around 13,000 contigs. The probes performed equally well in pools of the closely related species *Pinus sibirica*; in both species, more than 70% of the targeted contigs were sequenced at a depth ≥ 40x, i.e. the number of haplotypes in the pool. However, a thorough analysis of individually sequenced *P. cembra* samples indicated that a majority of the contigs (63%) represented multi-copy genes. We therefore removed paralogous contigs based on heterozygote excess and deviation from allele balance. Without putatively paralogous contigs, allele frequencies of population pools represented accurate estimates of individually determined allele frequencies. Using population genetic and landscape genomic methods, we show that inferences of neutral and adaptive genetic variation may be biased when not accounting for such multi-copy genes. Future studies should therefore put more emphasis on identifying paralogous loci, which will be facilitated by the establishment of additional high-quality reference genomes.

## Introduction

Investigating the genetic basis of local adaptation has recently received enormous attention in the field of evolutionary genetics and evolutionary ecology (reviewed in e.g., Flood & Hancock 2017; Tiffin & Ross-Ibarra 2014). It is now widely accepted that genetic signatures of natural selection are ideally studied with a (near to full) genome-wide approach (Hoban *et al.* 2016). Maximizing the investigated genomic space has several advantages. First, it decreases the probability of missing important genes or gene regions involved in local adaptation. Second, genome-scan methods like genome-wide association studies (GWAS, Korte & Farlow 2013), environmental association analyses (EAA, Rellstab *et al.* 2015), and population genomic methods like F_ST_ outlier or gene diversity analyses (Hohenlohe *et al.* 2010) often require characterizing the neutral population genetic structure of the studied populations to find (adaptive) regions in the genome that behave differently. Finally, the genetic basis of local adaptation is, in most cases, likely of polygenic nature (Csillery *et al.* 2018; Pritchard & Di Rienzo 2010), making it necessary to study gene interactions and pathways. Due to all these reasons, it is desirable to gather information of an as large as possible proportion of the genome.

Whole-genome re-sequencing has given interesting insights about how selection has left its traces in the genome, but such studies are often restricted to model organisms that have rather small and high-quality reference genomes (e.g., Exposito-Alonso *et al.* 2018; Günther *et al.* 2016; Machado *et al.* 2016) or a reference genome of a closely related species (e.g., Fischer *et al.* 2013). Despite the decreasing costs of sequencing in the past (https://www.genome.gov/sequencingcostsdata), projects with limited resources and/or study species with large genomes cannot afford re-sequencing of numerous individuals in several populations, as it is needed in population genomic studies. Under such conditions, other genotyping approaches have to be explored. These include, for example, targeted sequencing of candidate genes (e.g., Rellstab *et al.* 2016), SNP arrays (e.g., Barth *et al.* 2017), restriction site-associated sequencing (RAD-Seq) and similar approaches (e.g., Laporte *et al.* 2016), RNA sequencing (RNA-Seq, e.g., De Wit & Palumbi 2013) or exome capture (e.g., Yeaman *et al.* 2016). Moreover, the sequencing or genotyping of pooled DNA (Pool-Seq, normally at the population level) allows for reduced costs per individual and thus increased sample size (e.g., Waldvogel *et al.* 2017). All these approaches have their pros and cons in respect to the study of local adaptation, which are reviewed elsewhere (e.g., Hoban *et al.* 2016; Rellstab *et al.* 2015; Schlötterer *et al.* 2014).

From the approaches mentioned above, exome capture (Bamshad *et al.* 2011) represents one of the most interesting alternatives to whole-genome sequencing. By hybridizing to specifically designed probes, this approach targets the coding regions of the genome and consequently requires a priori knowledge of the coding sequences. In the absence of genomic resources, this information is best acquired by RNA-Seq and subsequent transcriptome assembly. Exome capture has been successfully applied in non-model species with large genomes, some of them over 20 Gbp (e.g., Neves *et al.* 2013; Suren *et al.* 2016; Yeaman *et al.* 2016), and is a promising approach for population-based genome scans and association studies (Jones & Good 2016). Disadvantages of using exome capture include the fact that non-coding regions are ignored (including introns that can interfere with hybridization), the labor-intensive design of probes (but see Puritz & Lotterhos in press), and the large variation in capture efficiency among probes and samples (Bamshad *et al.* 2011; Neves *et al.* 2013).

As holds for most conifers (Pinophyta), the genus *Pinus* (Pinaceae) is known for its species having large and complex genomes with estimated sizes that vary between 16 and 35 Gbp (Murray *et al.* 2012). Consequently, a reference genome for any pine species has been lacking until recently, and genomic studies mostly concentrated on characterizing transcriptomes (e.g., Pinosio *et al.* 2014) or genotyping populations using SNP arrays (e.g., Eckert *et al.* 2010; Mosca *et al.* 2016). However, in the last years, draft genome assemblies of the north-American species *Pinus taeda* (Wegrzyn *et al.* 2014; Zimin *et al.* 2014) and *P. lambertiana* (Stevens *et al.* 2016) were published. There is evidence for a whole-genome duplication event (WGD, with subsequent re-diploidization) preceding the Pinaceae radiation (Li *et al.* 2015). Yet, this WGD seems to a minor part responsible for the large genome sizes of *Pinus* species, which is mainly attributed to the high frequency of retrotransposons (Morse *et al.* 2009). The assemblies of *P. taeda* and *P. lambertiana* comprise 74 and 79% of such transposable elements (TEs), respectively. In addition, there is evidence for gene duplication, pseudogenes, and paralogs (Wegrzyn *et al.* 2014 and references therein). The immense size, complexity, and repetitive nature of pine genomes therefore represent a big challenge for population-based studies.

The existence of repetitive elements (hereafter referred to as "copies" or "paralogs") is a major issue for studies dealing with allele frequencies. The copies are similar enough to be mistaken as the same sequence during lab processes (binding of primers, hybridization of probes) and bioinformatic analyses (assembly or mapping), but contain private alleles that lead to false allele frequency estimates. However, filters and methods have become available to exclude such loci post-hoc (reviewed in McKinney *et al.* 2017; Willis *et al.* 2017). These approaches refer to, e.g., upper threshold for read depth, deviation from Hardy-Weinberg equilibrium (HWE, excess of heterozygosity), deviation from usual read ratios (allele balance), reference genome alignment, and haplotyping. Not all methods and filters are applicable in every genotyping/sequencing approach. For example, read depth is often not uniform, especially in approaches like exome capture or RAD-Seq, and high-quality reference genomes are often not available.

In this study, we used RNA-Seq to assemble the transcriptome of the conifer *Pinus cembra*, a tree species with an extremely large genome (estimated size of 29.3 Gbp, Zonneveld 2012) and no published reference genome. We show how RNA-Seq of different tissues, life stages, individuals, and treatments enlarges the sampled gene space as compared to using a single library only, and use the assembled transcriptome to design probes for exome capture of pooled population samples. The use of DNA pools instead of individuals in exome capture has shown to deliver accurate allele frequency estimates, even of rare variants (Bansal *et al.* 2011; Ryu *et al.* 2018). We demonstrate the importance of identifying and removing putatively paralogous loci, because they may have severe consequences on both the characterization of neutral and adaptive genetic variation. Finally, we show that our probes can also be successfully applied to a closely related species, *P. sibirica*. With this approach, we demonstrate possible avenues and potential challenges in genomic studies of non-model species with large genomes and a high proportion of paralogous genes.

## Materials and Methods

### Study species

*Pinus cembra* L. is a five-needle pine of the subgenus *Strobus* (Gernandt *et al.* 2005). It is a keystone species of the timberline ecotone, occurring at high elevation (1500-2400 m) in the Central European Alps and in the Carpathian Mountains. The species is mostly outcrossing (Salzer & Gugerli 2012) and shows high levels of gene flow due to wind pollination. However, it exhibits limited dispersal by seed (Salzer 2011) due to its dependency on its main vector, the spotted nutcracker (*Nucifraga caryocatactes*). Also other biotic factors (e.g., understory vegetation) and climatic factors seem to play an important role in post-dispersal recruitment (Meier *et al.* 2010; Neuschulz *et al.* 2018).

### RNA sequencing

To assemble the transcriptome of *P. cembra*, we performed RNA-Seq of different tissues, life stages, individuals, and treatments in winter and spring 2015. First, three juvenile plants (J1-J3) were exposed to different treatments. These plants were germinated in 2008 in the experimental garden of the Swiss Federal Research Institute WSL (Birmensdorf, Switzerland) using seeds from two different Swiss populations (Neuenalp for J1, Rautialp for J2 and J3, see Fig. 1 and Salzer & Gugerli 2012). For all treatments, fresh bud, needle, cambium, and root tissues were sampled if possible. First, we kept the plants under optimal growing conditions (22°C during day, 15°C at night, 16 hours of daylight, regular watering) in the greenhouse for 49 days to sample the control (C) treatment. Then we moved the plants outside and exposed them to freezing temperatures for one night (−5°C) to collect the frost-treated (F) samples, with around half a day acclimation at intermediate temperature before and after the treatment. Finally, we moved the juveniles back to the greenhouse and applied a drought treatment (D) during which they were exposed to control conditions, except that watering was extremely limited (once in 28 days). After this period, the plants showed symptoms of drought stress and were sampled. Second, we sampled a whole seedling (S, grown from a seed originating from Grächen, Switzerland, Fig. 1) that was kept under similar conditions as described above. Third, we sampled needles, cambium, and female and male flowers from an adult (A) tree planted in the WSL arboretum, presumably originating from south-eastern Switzerland.

**Figure 1:**
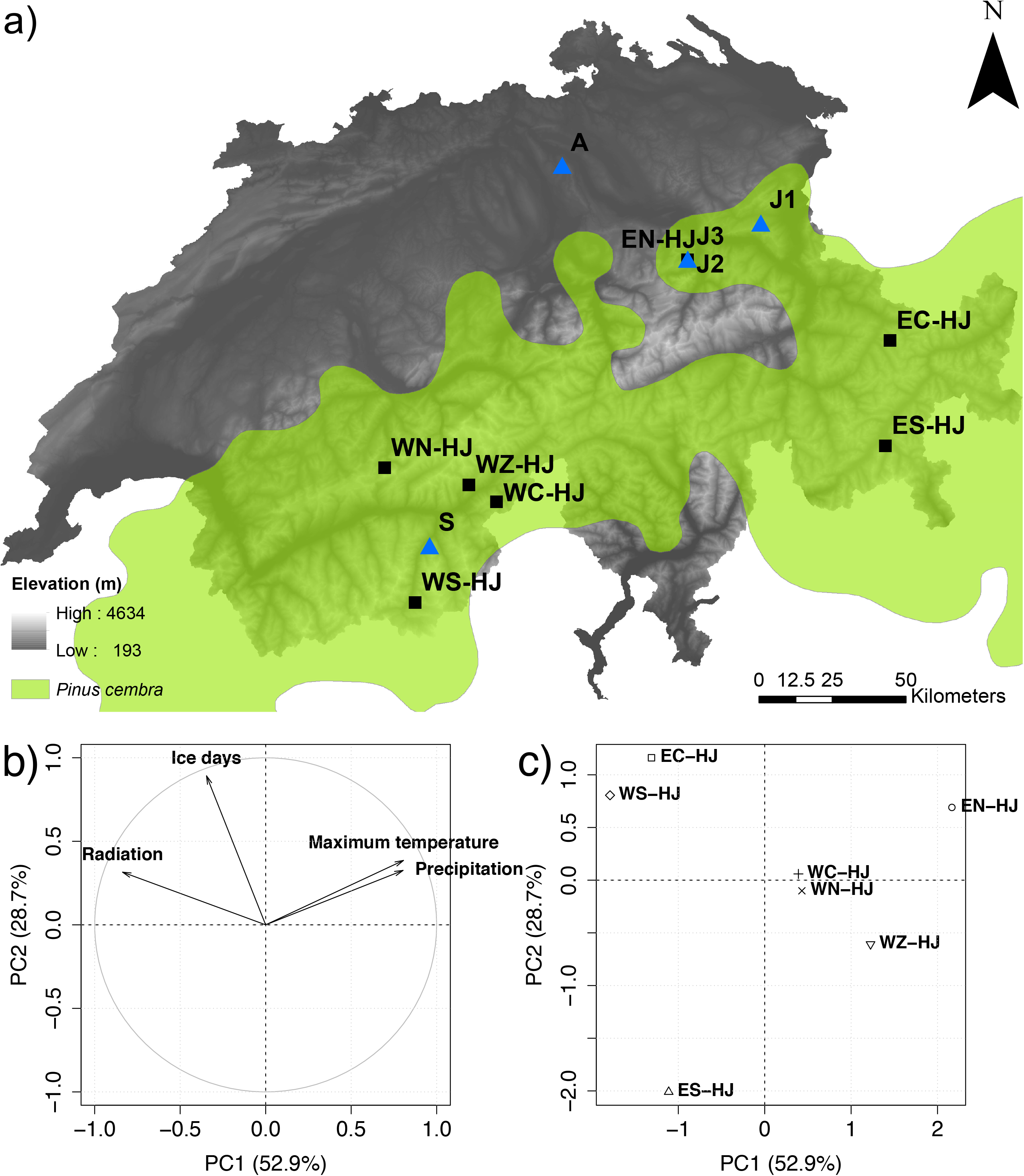
Description of sampling sites. (a) *Pinus cembra* distribution (green, from Caudullo *et al*. 2017) and localities of populations from where samples were taken for generating the transcriptome (triangles) and for exome capture (squares). Principal components analysis of environmental factors (b) and habitat characterization for the populations used in population genetic and environmental association analyses (c). Note that the adult tree (A) grows outside of its natural range and most likely originates from south-eastern Switzerland.

All samples were immediately transferred to liquid nitrogen and then kept at −80°C until further processing. RNA was extracted using the Plant RNA Isolation Mini Kit (Agilent, Santa Clara, USA). Quality and quantity of the RNA was checked with a Bioanalyzer (Agilent), a Quantus Fluorometer (Promega, Fitchburg, USA), and a Nanodrop (ThermoFisher, Waltham, USA). Per individual and treatment, we equimolarly pooled RNAs of all tissues, leading to eleven pools for sequencing (Table S1). Not all pools contained all tissues because repeated RNA extraction had failed in some cases. Libraries were prepared with the TruSeq RNA Library Kit (Illumina, San Diego, USA) and sequenced (paired-end reads of 125 bp) on two lanes of a HiSeq 2500 (Illumina). Library preparation and sequencing was performed by the Functional Genomics Centre Zurich (FGCZ), Switzerland.

### Transcriptome assembly and annotation

Raw reads were quality-trimmed with TRIMMOMATIC 0.35 (Bolger *et al.* 2014) using a quality threshold of 15 along a sliding window of 5 bp. The residual reads were then pooled for each individual and duplicate reads were removed with filterPCRdupl.pl (https://code.google.com/archive/p/condetri/downloads) using the first 100 bases from each read for comparison. We first assembled the transcriptomes of the five trees individually (ignoring treatments if there were any) using TRINITY (Grabherr *et al.* 2011) with a minimum transcript length of 200 bp and no Jaccard clipping. We then chose the assembly with the highest number of transcripts as the base assembly. The transcripts of the other four assemblies that were not part of the base assembly were added later (see Table S2 for details). The final assembly only included transcripts with a partial or complete open reading frame (ORF) and a plant BLAST hit on the NCBI Plant RefSeq database (https://www.ncbi.nlm.nih.gov/refseq) or Pfam (http://pfam.xfam.org). Low-coverage transcripts, ribosomal RNA, and contigs with signs of contamination (BLAST hit only with fungi, human, bacteria in the NCBI Nucleotide database [NCBI nt, https://www.ncbi.nlm.nih.gov/nucleotide]) were discarded. Finally, we kept only transcripts with a top BLAST hit on plants (NCBI nt). Functional annotation of the final assembly was done by BLASTing to NCBI nt, adding INTERPROSCAN (Jones *et al.* 2014) results, and processing in BLAST2GO (Götz *et al.* 2008). To analyze which transcripts (genes) were expressed in the different libraries (i.e., life stages, individuals, and treatments), we performed a simple presence/absence gene expression analysis (Appendix 1).

### Probe design for exome capture

We used the MYcroarray myBaits custom capture approach (Arbor Biosciences, Ann Arbor, USA). Probe design was done in-house using a series of perl and shell scripts (for details, see Table S3). To start, we only used the transcript of a “gene” defined by TRINITY with the highest expression. For these sequences, we determined the start and stop positions of the ORF prediction and extracted the coding sequences of each transcript. On these sequences, we placed 150-bp probe bases, always consisting of two 100-bp probes, with an overlap of 50 bp. We chose one probe base for sequences <2000 bp, two probe bases for sequences between 2000 and 4500 bp, and 3 probe bases for longer sequences. For TRINITY genes without probe base so far, we applied a second probe design approach as follows. Artificial reads of 100 bp in length and starting every 10 bp were generated from the transcript sequences. These reads were then mapped back to the transcripts with BWA MEM. Any sequence position in the transcripts with secondary or supplementary read mappings or coverage either zero or higher than 10 was masked. For all transcripts without probes in the initial approach we extracted all stretches of at least 100 bp from the unmasked regions. On these potential probe bases we placed tiled probes as in the initial approach. The probes from the two approaches were combined and filtered for sequence similarity (> 90%) and overlap (> 50%). We also removed probes from highly similar genes (≥ 98% cluster identity). Using DUSTMASKER (Morgulis *et al.* 2006), we identified probes that contained repetitive regions longer than 40 bp, such as microsatellites or low-complexity regions and removed them. Finally, we removed probes with a GC content ≥ 70%.

### Population sampling, DNA extraction, and exome capture

In summer 2014, we sampled needles of 25 georeferenced trees in seven *P. cembra* populations covering a wide range of the Swiss Alps (Fig. 1, Table S4). We concentrated on juvenile trees (estimated 7-20 years based on annual shoot increment) above the timberline. DNA of 15-20 mg lyophilized needle material was extracted using the DNeasy 96 Plant Kit (Qiagen, Hilden, Germany). DNA quality was checked on an agarose gel containing 1% agarose and 1x TBE, and with a Nanodrop. DNA quantity was measured with a Quantus Fluorometer. For each population, we selected 20 trees based on DNA quality and quantity of the extractions to produce equimolar DNA pools consisting of 55 ng DNA of every individual (in total 1100 ng).

For the closely related *P. sibirica*, we used DNA of seeds originating from East Siberia (Krasnoyarsk, Russia). Seeds were collected from three different forest districts far from each other (250-500 km), namely Yeniseiskoye (sample PS01), Kozulskoye (PS04), and Tashtynskoye (PS06). We extracted DNAs of 20 lyophilized embryos per district as described above and created one equimolar DNA pool per district. Note that these samples do not represent populations, but a small subsample of large commercial seed collections from large forest districts.

We sheared the DNA pools with a Q800R2 Sonicator (Qsonica, Newtown, USA) for 10 min and created barcoded libraries (average size of 550 bp measured with an Agilent 2200 Tape Station) using the NEBNext Ultra II DNA Library Prep Kit for Illumina (New England Biolabs, Ipswich, USA). To compare allele frequencies derived from pooled vs individual sequencing, we also created libraries of the individuals included in population pools EC-HJ and WC-HJ. After library preparation, hybridization of probes was performed using the MYcroarray myBaits custom capture kit according to the manufacturer’s instructions, including an amplification step with 14 cycles. The 50 hybridized libraries (seven population pools and 2x 20 individuals of *P. cembra*, three pooled samples of *P. sibirica*) were then super-pooled for sequencing on a total of four lanes of an Illumina HiSeq 4000 (paired-end reads of 150 bp). Sequencing was performed by FGCZ and Fasteris (Geneva, Switzerland). Because pooled sequencing requires higher coverage than individual sequencing, the super-pools contained ten times more material from the pools than from individuals. The obtained reads were filtered and trimmed with TRIMMOMATIC for quality 20 along a sliding window of 5 bp. Only read pairs for which both forward and reverse reads passed the quality criterion were further considered. Reads were then mapped with BOWTIE 2.3.0 (Langmead *et al.* 2009) to those contigs of the transcriptome that contained probe bases (options -p 4 -I 0 -X 540 -D 25 -R 5 -N 0 -L 18 --gbar 8 --mp 8,3 -i S,1,0.30). SNP calling was performed with GATK (McKenna *et al.* 2010) with ploidy set to 2 for individuals and 40 for pools.

### Identification and filtering of putatively paralogous genes

We first called SNPs for the 40 individual samples and the corresponding population pools (EC-HJ and WC-HJ) to identify putatively paralogous contigs. We kept SNPs with coverage ≥ 8x (individuals) or ≥ 80x (pools) and mapping quality/depth ratio ≥ 0.25.

To identify putatively paralogous contigs, we used the HDPLOT approach of McKinney *et al.* (2017) with individual genotype data. For every locus, this method assesses both the frequency of heterozygous individuals (*H*) and the deviation (*D*) from expected read ratio (RR) in heterozygous individuals, and then uses specific thresholds for *H* and *D* to identify putatively paralogous loci. Using the case of the Chinook salmon (*Oncorhynchus tshawytscha*) reference genome, where single-copy and paralogous gene regions are exactly known, McKinney *et al.* (2017) showed that this method reliably identifies paralogous loci. In single-copy genes, *H* should not exceed 0.5 under HWE and RR should be around 1. Loci showing signals of selection are not expected to be in HWE, however, in a scenario of divergent selection where beneficial alleles have the tendency to move towards fixation in certain environments, one would expect reduced heterozygosity. Setting an upper threshold for *H* therefore does not exclude putatively adaptive (divergent) loci from further analyses. RR represents the ratio between the number of reference and alternative allele reads in heterozygous individuals (also called allele balance). *D* results from a statistic describing RR that takes sequencing depth into account and reduces (compared to raw RR values) the probability of misclassifying single-copy loci as multi-copy loci at low minor allele frequencies (MAF). HDPLOT was run in R 3.4.4 (R Development Core Team 2018) using the code available at McKinney *et al*. (2016) with the vcf files as input data.

Thresholds for *H* and *D* were set after visual inspection. Every SNP in each individual was either set to putatively single-copy (no threshold exceeded) or multi-copy (one of the two thresholds exceeded) in both individually genotyped populations. Contigs with at least two multi-copy SNPs (or one, if max. two SNPs were present) in at least one population were considered as multi-copy. The effectiveness of filtering paralogous contigs was assessed by looking at the correlation between allele frequencies derived from individual vs pooled sequencing and the allele frequency spectra (AFS) of both individuals and pools before and after filtering. AFS of pooled samples are much less sensitive to an excess of *H* and *D*, because the pooled allele frequencies are not subjected to a genotyping step, but represent the ratio of read counts. We also checked the effect of filtering for several other parameters that are indicated in the vcf files; SNP quality (Q), read depth (RD), difference in mapping qualities of reads supporting the reference vs alternative allele (mapping quality rank sum, MQRS), strand odds ratio to detect allele specific strand bias (SOR), and position bias of reference and alternative alleles (read position rank sum, RPRS).

### Population genetic structure and environmental association analysis using the full and single-copy datasets

Next, we wanted to know if using all SNPs (hereafter referred to as the "full dataset") shows different patterns in respect to neutral and adaptive genetic variation than when only using SNPs of single-copy genes (hereafter referred to as "single-copy dataset"). We called SNPs for all seven population pools and kept SNPs with coverage ≥ 40x and mapping quality/depth ratio ≥ 0.25. We only used SNPs without missing data and MAF of ≥ 2.5% in at least one population.

We applied a series of multivariate and population genetic analyses to characterize the neutral genetic variation among and within populations for both the full and the single-copy datasets. First, we performed a principal component analysis (PCA) using the "prcomp" function in R. We then estimated the similarity of the first two principal components (PCs) of both datasets using a Spearman correlation of the respective principal coordinates in R. Furthermore, we estimated pairwise *F*_ST_ (Weir & Cockerham 1984) between populations and global *F*_ST_ using the "calcPopDiff" function in the R package POLYSAT (Clark & Jasieniuk 2011) with 1000 bootstrap replicates and ploidy set to 40 that corresponds to the number of haplotypes in the pool. The correlation of pairwise *F*_ST_ of both datasets was tested with a Mantel test (Mantel 1967) based on 10,000 permutations using the R package VEGAN (Oksanen *et al.* 2013). Finally, we compared genetic diversity within populations among the two datasets by calculating expected heterozygosity *H*_e_ as in Fischer *et al.* (2017) and the proportion of polymorphic loci (PPL). Potential differences in *H*_e_ and PPL between the two datasets were tested with a Pearson’s correlation and a paired *t* test in R.

Environmental association analysis (EAA) correlates allele frequencies and environmental descriptors of the sampling locations to identify loci putatively involved in adaptation to the local environment (Rellstab *et al.* 2015). For this, we extracted four topo-climatic factors (number of ice days, maximum temperature, precipitation, and solar radiation; Tables S4 and S5) for every georeferenced individual and retained the average of each population for EAA. The four environmental factors were not highly correlated (Pearson’s *r* < 0.7; Table S6).

In the EAA, we tested for linear associations between environmental factors and allele frequencies with latent factor mixed models (LFMM, Frichot *et al.* 2013). LFMM has shown to be a robust method for detecting candidate loci potentially under selection (de Villemereuil *et al.* 2014; Lotterhos & Whitlock 2015) by incorporating the neutral genetic structure (as random latent factors) in combination with rigorous tests to control for false-positive associations (François *et al.* 2016). We used LFMM 2.0 incorporated in the R package LFMM (Caye & François 2018). LFMM can handle population allele frequencies, but does not account for the pool size. As in the BAYENV approach of Günther and Coop (2013), we therefore artificially increased our sample size by resampling 20 individual allele frequencies for each pool at each locus from a beta distribution (Waldvogel *et al.* 2017). Then, we carried out the EAA between the simulated individual allele frequencies and the population-based environmental factors. We used population averages of environmental factors, because individual environment values cannot be linked to randomly sampled individual allele frequencies. We ran the function "lfmm_ridge" for both the single-copy and the full datasets with 20 repetitions (including resampling of individual frequencies) for K = 1-7 for each environmental factor, and calculated the genomic inflation factor (λ) from median *z* scores with the function "lfmm_test". We adjusted the *p* values based on λ and a χ^2^ distribution (François *et al.* 2016). We then applied the Benjamini-Hochberg algorithm (Benjamini & Hochberg 1995) to identify significant associations with q < 0.001. To visualize the proportion of associated SNPs shared between both datasets, we used the R package VENNDIAGRAM (Chen & Boutros 2011) for each environmental factor with *K* = 2 and 4. We then carried out Pearson’s correlations between the median z scores of both datasets exemplarily for *K* = 2.

### Comparison between *P. cembra* and *P. sibirica*

To verify whether our probes designed for *P. cembra* also efficiently hybridized in *P. sibirica*, we assessed the number of contigs that contained at least one position with a coverage ≥ 40x (the threshold used for SNP calling) with the "genomecov "function of BEDTOOLS (Quinlan & Hall 2010) and the number of SNPs before and after filtering (MAF ≥ 2.5% in at least one sample, no missing data).

## Results

### RNA sequencing and transcriptome assembly and annotation

The Illumina sequencing of the eleven RNA libraries resulted in 498.3 million read pairs. Of those, 96.9% passed the quality trimming step and 53.5% remained after removing duplicate reads, leading to 266.8 million read pairs for the transcriptome assembly (Table S1). The highest number of trimmed and de-duplicated read pairs was available for sample J1 (90.5 million) and the smallest number for S (21.3 million).

The five initial assemblies, including reads of all treatments, led to a range of 148,234 (A) to 403,403 (J1) transcripts per assembly. Based on this number and other parameters (read length distribution, read content, E90-N50 statistics), we chose J1 as the base assembly. The final transcriptome consisted of 40,468 transcripts representing 27,979 TRINITY genes, totaling to 69.4 Mbp of sequence. Average and median transcript length were 1711 and 1422 bp, respectively (Fig. S1). N50 was 2380 bp and GC content 42.2%. More than 80% of the contigs (34,010) could be annotated. Most BLAST top-hits were on genes of the conifer *Picea sitchensis*, followed by two angiosperm species (Table S7). With *Pinus taeda* and *P. tabuliformis*, two pine species were also in the top ten of the top-hit species list. In total, 12,038 different proteins were present in the annotation, of which 5656 were annotated uniquely to a transcript. The GO terms assigned to the annotated proteins were dominated by metabolic processes, but functions in relation to the applied treatments before RNA-Seq were also prominently represented (Table 1).

**Table 1:** Characteristics of the Pinus cembra transcriptome. Top 20 gene ontology (GO) terms (only biological processes) of the annotated transcripts of the transcriptome assembly, and GO terms with > 100 hits related to the treatments applied to the three juvenile samples before RNA-Seq.

| Rank | GO-term | Description | No. of proteins |
| --- | --- | --- | --- |
| 1 | GO:0055114 | oxidation-reduction process | 2847 |
| 2 | GO:0006355 | regulation of transcription, DNA-dependent | 2483 |
| 3 | GO:0008152 | metabolic process | 2288 |
| 4 | GO:0006468 | protein phosphorylation | 1564 |
| 5 | GO:0009069 | serine family amino acid metabolic process | 1359 |
| 6 | GO:0006508 | proteolysis | 863 |
| 7 | GO:0006118 | electron transport | 813 |
| 8 | GO:0055085 | transmembrane transport | 667 |
| 9 | GO:0007165 | signal transduction | 653 |
| 10 | GO:0005985 | sucrose metabolic process | 641 |
| 11 | GO:0005982 | starch metabolic process | 576 |
| 12 | GO:0005975 | carbohydrate metabolic process | 516 |
| 13 | GO:0042254 | ribosome biogenesis | 508 |
| 14 | GO:0006412 | translation | 496 |
| 15 | GO:0006952 | defense response | 483 |
| 16 | GO:0006094 | gluconeogenesis | 474 |
| 17 | GO:0044763 | single-organism cellular process | 466 |
| 18 | GO:0006096 | glycolysis | 452 |
| 19 | GO:0016310 | phosphorylation | 418 |
| 20 | GO:0006810 | transport | 372 |
| 21 | GO:0009651 | response to salt stress | 355 |
| 23 | GO:0006979 | response to oxidative stress | 342 |
| 24 | GO:0006950 | response to stress | 339 |
| 55 | GO:0009737 | response to abscisic acid stimulus | 193 |
| 59 | GO:0009409 | response to cold | 182 |
| 64 | GO:0009408 | response to heat | 172 |
| 81 | GO:0009414 | response to water deprivation | 153 |
| 131 | GO:0009266 | response to temperature stimulus | 115 |

### Probe design, population sampling, and exome capture

In total, we designed 54,870 probes for 24,998 transcripts of the transcriptome assembly. Of these, 22,766 transcripts contained two (tiled) probes, 2,027 transcripts four probes, and 205 transcripts six probes. The sequencing yielded 1.646 billion read pairs from the exome capture of the seven population pools, 40 individual trees, and three *P. sibirica* pools (Table S8). After adapter and quality trimming, 94.3% of these read pairs remained. In the 50 libraries, 58.3 to 78.1% of the original read pairs mapped back to the targets with a minimum mapping quality of 20. Average mapping success was higher in the individuals of EC-HJ (75.5%) than in those of WC-HJ (64.0%). In the *P. cembra* pools, an average of 65.1% of the original read pairs mapped back to the targets, whereas it was 61.3% for *P. sibirica* samples.

### Genotyping and filtering for paralogs

To identify putatively paralogous contigs, we concentrated on populations EN-HJ and WC-HJ, for which we had both individual and pooled sequences at hand. Results in this section present both populations (EN-HJ and WC-HJ) together; separate analyses for the two single populations led to highly similar outcomes (results not shown). After basic filtering (QD ≥ 0.25 and DP ≥ 80 for pools or DP ≥ 8 for individuals) and reducing the SNP list to those that were found both in individual and pooled datasets, we remained with 95,074 SNPs in 10,675 contigs (42.7% of the original transcripts) in both populations. We found 82,658 of those SNPs in 10,207 contigs in EC-HJ and 66,607 SNPs in 8888 contigs in WC-HJ. Of these, 53,661 SNPs and 8420 contigs were shared between the two tested populations.

The SNP set showed strong signs of paralogous genes (Figs. 2 and 3). Although population allele frequencies from pools and individuals were significantly correlated (Pearson’s *r* = 0.81, *p* < 0.001), both the density plot (Fig. 2a) comparing population allele frequencies from individuals and pools, and the AFS of the individual samples (Fig. 2b) showed a clear heterozygosity excess. This was not visible in the AFS of the pooled samples (Fig. 2c). Also the HD plot (Fig. 3a) revealed a clear heterozygosity excess (*H*, Fig. 3b) and a strong deviation from the expected allele ratio (*D*, Fig. 3c), visible by a second peak at around *D* = 7. The latter means that, in heterozygous individuals, reference and alternative alleles are not always supported by similar numbers of reads. Based on visual inspection of Fig. 3, we decided to classify SNPs with *H* ≥ 0.6 and *D* ≥|4| as multi-copy. After removing putatively paralogous contigs according to the rules defined in the Material and Methods section, we remained with 4950 contigs containing 14,499 SNPs. The correlation between allele frequencies from individual and pooled samples using these filtered contigs was very strong (Pearson’s *r* = 0.96, *p* < 0.001, Figure 2d), and the AFS of the individual samples was close to the expected L-shape (Figure 2e). We therefore considered these 4950 contigs as single-copy genes. Importantly, our analyses showed that in both tested populations EN-HJ and WC-HJ, roughly the same contigs were classified as single-copy or multi-copy (Table S9). Of 10,675 contigs, 7680 were assigned to the same category in both populations. Only 669 resulted in contrasting outcomes in the two populations and were therefore conservatively classified as potential multi-copy contigs according to the rules described above. The rest of the contigs did not contain data in at least one of the populations.

**Figure 2:**
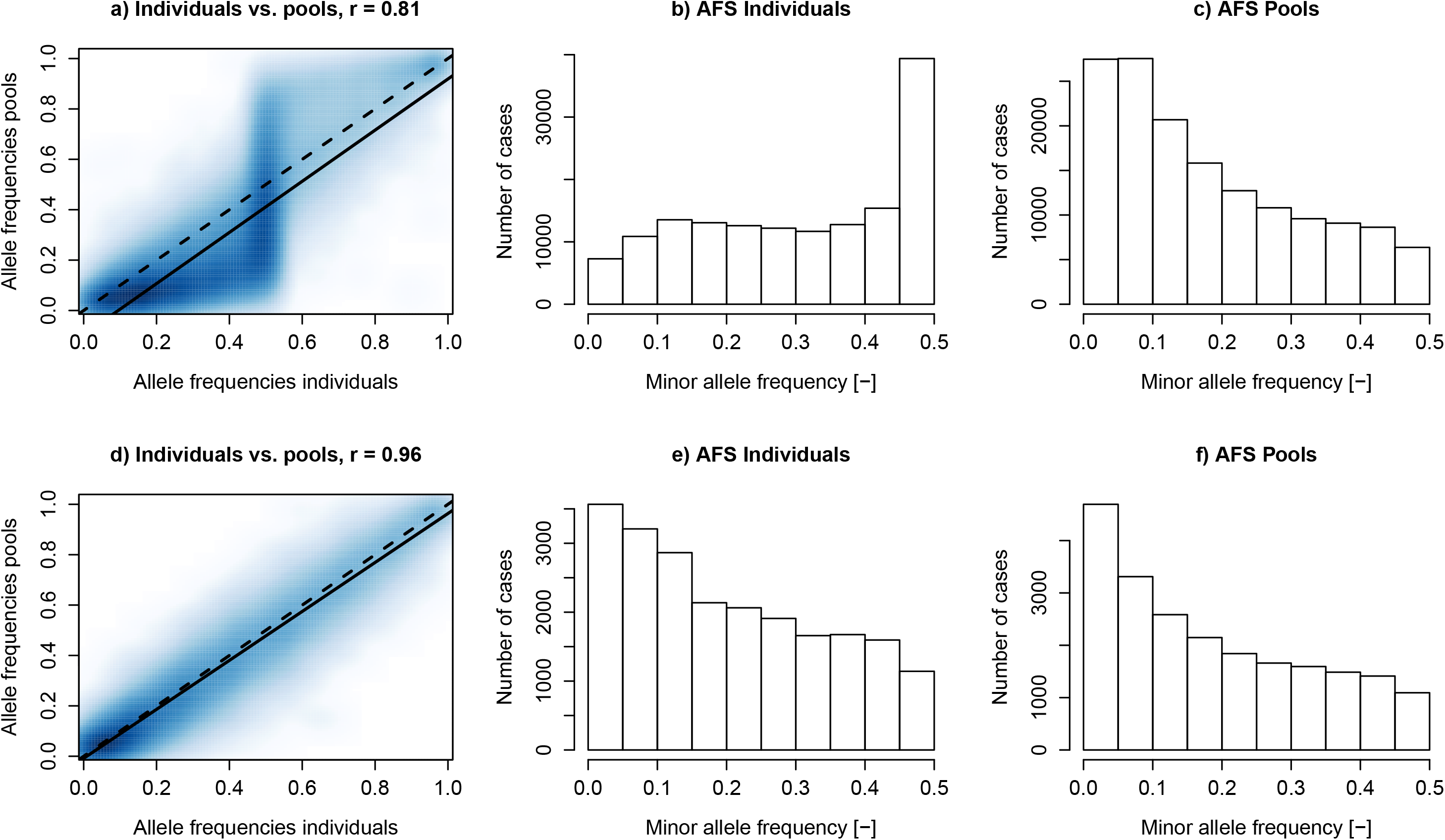
Characteristics of the obtained SNP set before and after filtering for putatively paralogous contigs in *Pinus cembra*. Plots represent the situation before (a-c) and after filtering (d-f) for excess heterozygosity and deviation from expected read ratio (see Fig. 3). Shown are density plots of the comparison between allele frequencies derived from individual and pooled samples including Pearson’s correlation coefficient *r* (a, d). The solid line represents a linear regression and the dashed line the expected 1:1 ratio. Shown is also the minor allele frequency distribution (allele frequency spectrum, AFS) of the individual (b, e) and pooled (e, f) samples. Data were combined from the two individually sequenced populations EN-HJ and WC-HJ.

**Figure 3:**
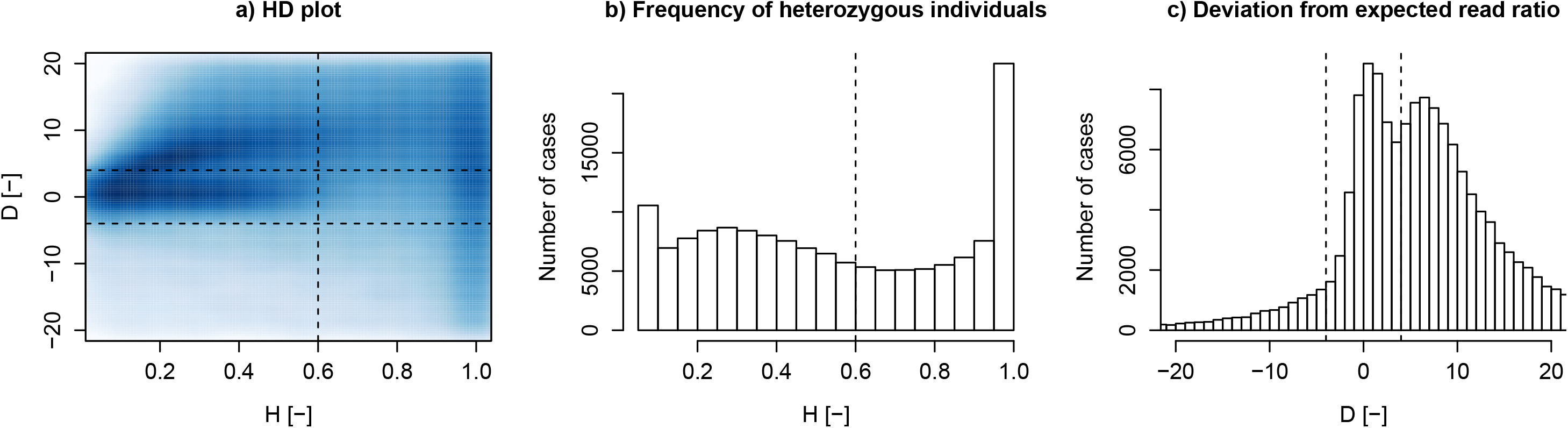
Applying the HD plot method of McKinney *et al.* (2017) in *Pinus cembra*. Characteristics of the obtained SNPs (density plot; a) before filtering for excess heterozygosity (*H*; b) and deviation from expected read ratio (*D*; c). Applied thresholds are depicted as dashed lines. Data were combined from the two individually sequenced populations EN-HJ and WC-HJ.

From the other five filters tested (Fig. S2), only the mapping quality rank sum (MQRS, Fig. S2e) improved the correlation between individual and pooled allele frequencies, and slightly changed the AFS. This indicates that the reads supporting the alternative alleles generally had lower mapping quality scores than those supporting the reference allele. Filtering using the remaining four parameters (*Q*, RD, SOR, or RPRS) did not change or even worsen the outcome. Specifically, read depth (RD, Fig. S2d), which could be indicative of paralogous genes (e.g., McKinney *et al.* 2017), did not show a multimodal distribution, and filtering for high RD did not improve results. Note that the scenarios in Fig. S2 only show one (reasonably, but arbitrarily) chosen threshold per parameter. Using other thresholds did not notably improve the correlation and AFS (data not shown).

### Population genetic structure and environmental association analysis using the full and single-copy datasets

After mapping and basic filtering, we obtained 221,735 raw SNPs in 16,264 contigs across all seven populations. From these SNPs, we discarded 103,810 SNPs because of missing data and 17,206 SNPs due to too low MAF. This resulted in a total of 100,719 SNPs for the full dataset, representing 10,220 contigs. For the single-copy dataset, we only retained those 3818 contigs that showed no signs of paralogs according to the HD plot analysis described above, comprising 13,328 SNPs.

Population genetic structure, analyzed with a PCA, was similar between the full and the single-copy datasets (Fig. 4). The first two PCs explained 38.7% (full dataset) and 37.2% (single-copy dataset) of the allele frequency variation among populations, respectively. Consistently, the first axis separated Eastern and Western populations, whereas the second axis revealed more substructure. The corresponding coordinates of both the first and second PC were correlated (Fig. 4c,d). Also pairwise FST values were highly correlated between the two datasets (Fig. S3a) and supported moderate differentiation among populations with significantly higher pairwise values for the single-copy dataset, ranging from 0.140 to 0.204 compared to 0.075-0.100 for the full dataset (Table S10). The two measures of global *F*_ST_ revealed the same tendency (Table S11). Both PPL and *H*_e_ of the two datasets were highly correlated, but significantly higher for the full than for the single-copy dataset (Table S11 and Fig. S3b).

**Figure 4:**
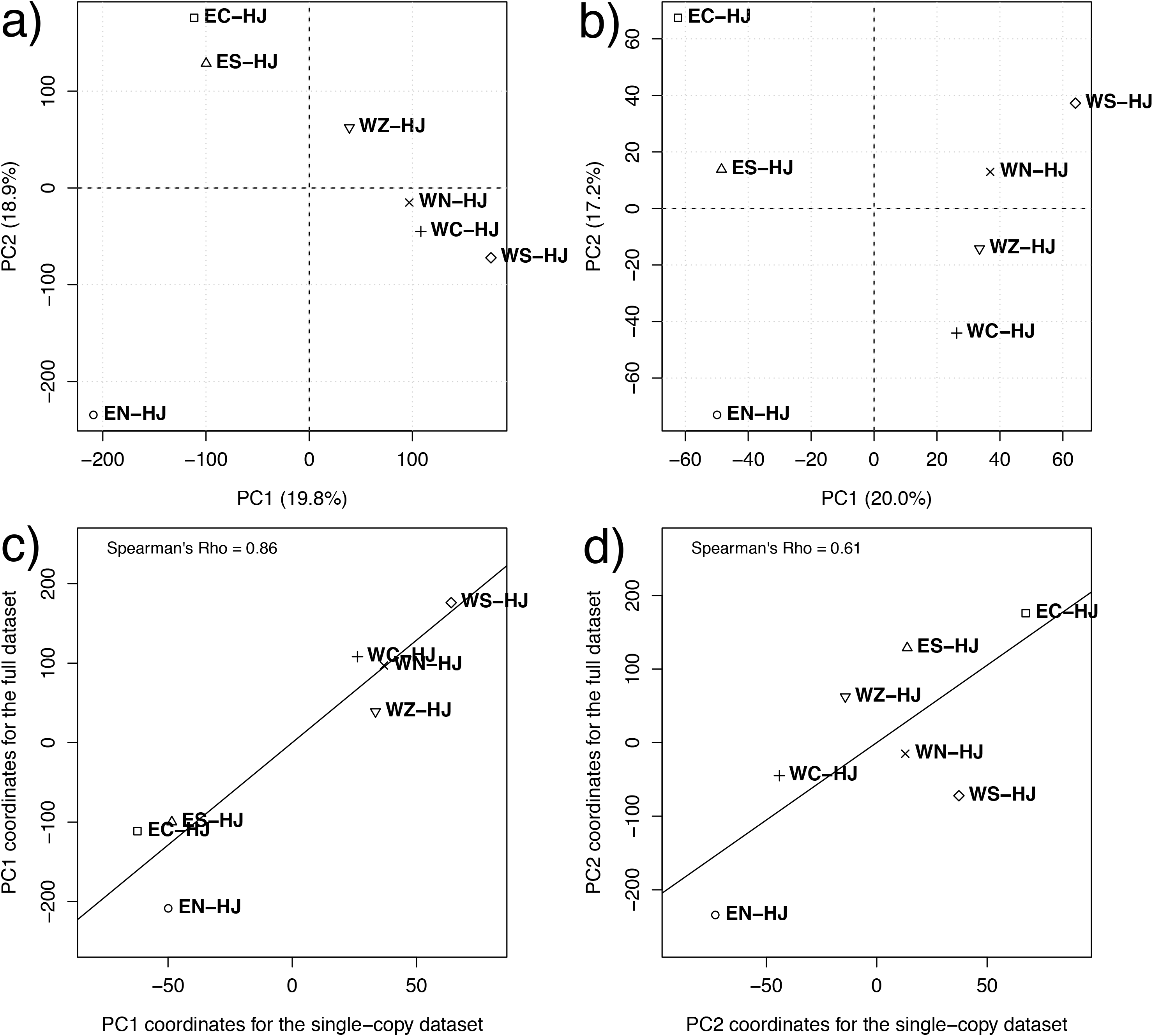
Analysis of neutral population structure using the full and single-copy datasets in *Pinus cembra*. Principal components (PC) analysis of pooled populations for the full (a) and the single-copy dataset (b). Correlation between the PC coordinates of both datasets for the first (c) and the second axis (d).

Despite the restricted altitudinal range of the populations, we found environmental variation among their habitats (Fig. 1c and Table S4). The two first PCs accounted for 81.6% of the variance and resulted in 4-5 environmental groups of populations irrespective of their geographic locations. The third axis roughly represented a West–East gradient (result not shown).

In the EAA, λ was always lower for the full dataset compared to the single-copy dataset, lowest for *K* = 1, and increased with increasing *K*. Depending on *K* and environmental factor, associated SNPs increased 8.8-35.0 fold for the full dataset compared to the single-copy dataset (Table S12). Due to the clear separation of Eastern and Western populations (suggesting *K* = 2) and the large increase in λ from *K* = 4 to 5 (Table S12, suggesting *K* = 4), we had a closer look at *K* = 2 and 4. For *K* = 2, we found 12,037 (full dataset) and 671 (single-copy dataset) associations of a SNP with one of the four environmental factors; for *K* = 4 it was 21,714 and 1667, respectively. For both *K* values, most of the associated SNPs identified in the single-copy dataset were also part of the associated SNPs in the full dataset (Fig. 5). The z scores of the shared associated SNPs of both datasets for *K* = 2 showed a significant positive correlation in all four environmental factors (Fig. S4).

**Figure 5:**
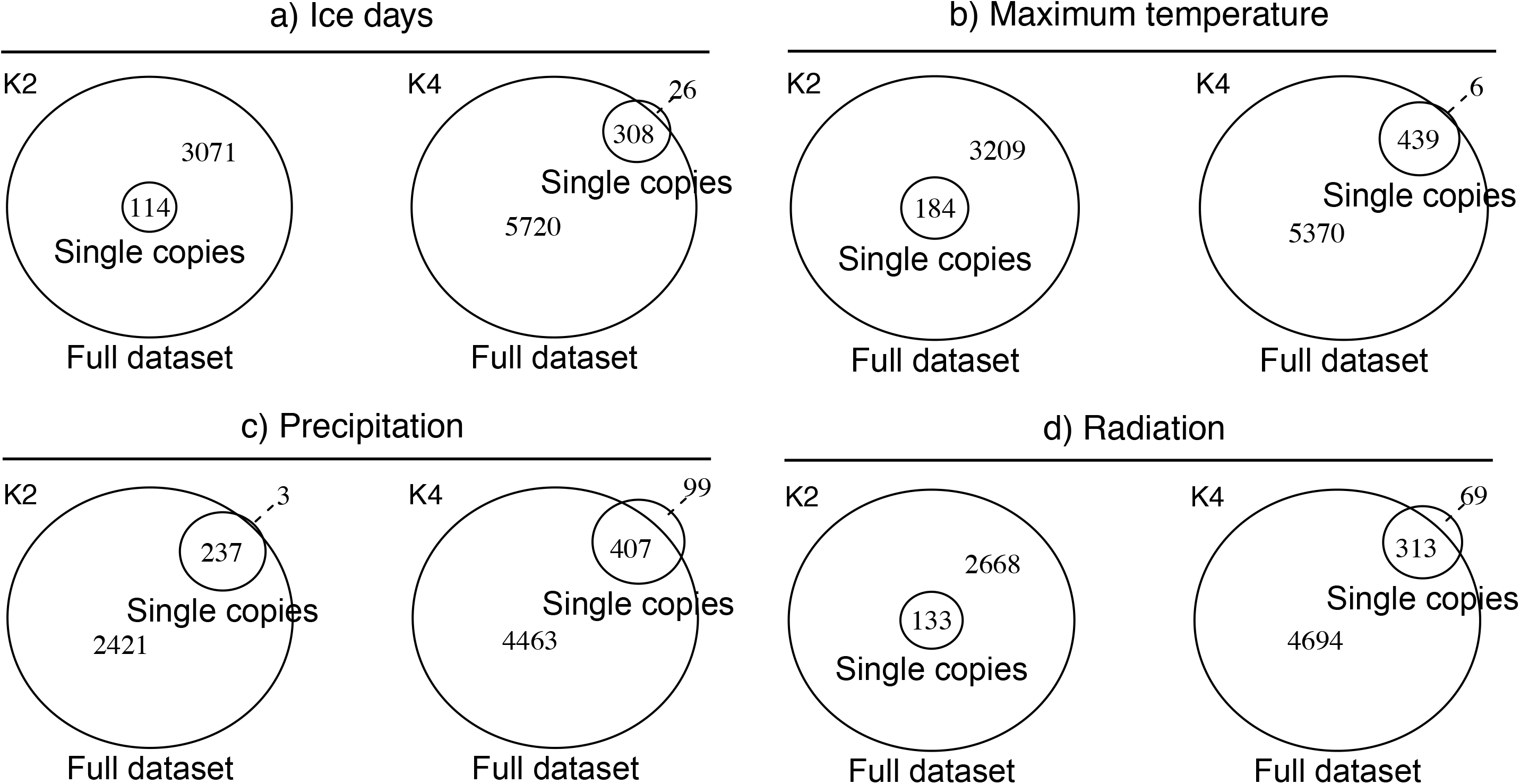
Overlap of SNPs associated to environmental factors between the full and single-copy datasets of *Pinus cembra*. Shown are the environmental factors ice days (a), maximum temperature (b), precipitation (c), and radiation (d) for both K = 2 and 4, the number of putative genetic clusters.

### Use of the probes in the closely related species P. sibirica

Of all mapped contigs, 81.9% (*P. cembra*) and 81.1% (*P. sibirica*) had a minimum coverage ≥ 40x in at least one of the positions in the contig (the required coverage threshold for the SNP calling). We found a similar number (for *p* values, see Table 2) of such contigs in the pooled *P. cembra* populations (average: 17,960 ± 575 SD) and the pooled *P. sibirica* samples (17,802 ± 381 SD). However, the P. sibirica contigs were significantly more polymorphic than those of P. cembra populations (Table 2). After SNP calling and filtering, we identified more SNPs in *P. sibirica* samples (107,810 SNPs ± 3822) compared to *P. cembra* populations (84,008 SNPs ± 7143), but the average number of SNPs in SNP-containing contigs did not differ significantly (8.7 ± 0.03 and 9.0 ± 0.44, respectively).

**Table 2:** Exome capture comparison for *Pinus cembra* and *P. sibirica.* Filtering means that we retained SNPs that exhibited a minor allele frequency of ≥ 2.5% in at least one population/sample and without missing data in all populations/samples. minRD = minimum read depth in at least one position of the contig.

| | Code | Contigs<br>(minRD $\geq 40x$) | Contigs with SNPs<br>(before filtering) | SNPs<br>(before filtering) | Contigs with SNPs<br>(after filtering) | SNPs<br>(after filtering) | SNP/contig<br>(after filtering) |
| --- | --- | --- | --- | --- | --- | --- | --- |
| *Pinus cembra* | EC-HJ | 17,807 | 12,802 | 140,182 | 9,335 | 88,931 | 9.5 |
|  | EN-HJ | 17,272 | 11,992 | 119,180 | 8066 | 70,416 | 8.7 |
|  | ES-HJ | 17,244 | 12,267 | 131,592 | 8,795 | 83,416 | 9.5 |
|  | WC-HJ | 17,987 | 12,755 | 128,244 | 9,219 | 79,399 | 8.6 |
|  | WN-HJ | 18,376 | 13,162 | 133,598 | 9,799 | 86,335 | 8.8 |
|  | WS-HJ | 18,817 | 13,550 | 136,526 | 10,433 | 88,446 | 8.5 |
|  | WZ-HJ | 18,214 | 13,154 | 140,716 | 9,759 | 91,111 | 9.3 |
|  | Average | 17,960 | 12,812 | 132,863 | 9,344 | 84,008 | 9.0 |
|  | Total | 19,427 | 14,896 | 166,336 | 13,152 | 141,566 | 10.8 |
| *Pinus sibirica* | PS01 | 17,886 | 15,081 | 157,675 | 12,548 | 108,486 | 8.6 |
|  | PS04 | 18,133 | 15,373 | 161,738 | 12,775 | 111,249 | 8.7 |
|  | PS06 | 17,386 | 14,609 | 153,495 | 11,917 | 103,695 | 8.7 |
|  | Average | 17,802 | 15,021 | 157,636 | 12,413 | 107,810 | 8.7 |
|  | Total | 18,605 | 16,286 | 176,497 | 15,000 | 152,986 | 10.2 |
| Student's t test | p value | 0.68 | < 0.001 | < 0.001 | < 0.001 | < 0.001 | 0.27 |
| Shared between species |  | 18,513 | 12,954 | — | 11,385 | — | — |

## Discussion

### RNA-Seq and transcriptome assembly

The newly established transcriptome for *P. cembra* is similar to other published pine transcriptomes (e.g., Baker *et al.* 2018; Wachowiak *et al.* 2015) in terms of total size (69.4 Mbp), number of transcripts (40,468), number of genes (27,979), median transcript length (1422 bp), and proportion of annotated transcripts (84%). Since our goal was to gain information for the largest possible gene space to design capture probes, we included multiple libraries consisting of various tissues of different life stages, individuals, and treatments. Our presence/absence-based gene expression analysis shows that this effort led to a substantial increase of characterized genes as compared to using a single source of biological material (Appendix 1). However, not all libraries equally added genes to the final assembly. Whereas including different life stages added the highest number and proportion of library-specific transcripts (Fig. A1.1b), the use of replicates (different juvenile individuals, Fig. A1.1c) and treatments (drought and cold stress, control; Fig. A1.1d) did not add many. However, this finding should be treated with caution, because we were not able to include all tissues in all libraries of the juvenile life stages (Table S1) and we cannot exclude the possibility that our treatments had a smaller effect than expected. Still, the relatively few genes expressed privately under drought and frost stress might actually represent interesting targets for studying adaptation to environmental variation. For example, the SUGAR TRANSPORT PROTEIN 13, only expressed in the drought treatment, plays an important role in the long-distance transport of sugar within the plant and has shown to be activated in drought and salinity stress in e.g. *Arabidopsis thaliana* (Yamada *et al.* 2011). On the other hand, the commonly expressed genes might represent genes that are always expressed, but at varying expression levels and/or in different variants (not assessed in our analysis). In conclusion, it seems to be most efficient to include different life stages rather than invest a lot of effort into treatments, but this comes with the cost of eventually missing important stress related genes.

### Pooled exome capture

Our pooled exome capture approach showed to be a powerful strategy to sequence coding regions of the genome. Between 58 and 78% of the raw reads could be mapped back to the targets (Table S8), and 72% of the targets contained positions with a coverage of at least 40x. This is higher than found in many other studies (for examples see Puritz & Lotterhos in press), which often differ in the type of reference that reads are mapped to. One possible reason for the existence of non-mapped reads are exon/intron boundaries located in the probe base region (Neves *et al.* 2013; Suren *et al.* 2016), which could be circumvented when a high-quality reference genome is available. It is important to emphasize that in this study, we do not always cover the whole sequence of the original transcript, but gain sequence information for one to three stretches (number of probe bases) typically summing up to 200-1500 bp (average = 861 bp, median = 764 bp, Figure S5).

The pooled allele frequencies proved to be a good proxy for the observed allele frequencies based on individual sequencing in populations EN-HJ and WC-HJ, at least after removing putative paralogous contigs (Fig. 2d). This is in line with other studies that used pooled DNA samples for exome capture (Bansal *et al.* 2011; Ryu *et al.* 2018). However, compared to other studies that compared pooled and individually genotyped populations (reviewed in Rellstab *et al.* 2013), the accuracy of the pooled allele frequency seemed to be slightly lower. The reason for this might be the more complex library construction in exome capture compared to other sequencing and genotyping approaches that normally do not include a hybridization step. Still, equimolarly pooling the DNA of populations is a highly cost-effective approach for population genetic analyses, with the disadvantage of losing individual genotype information.

Most importantly, our approach led to the identification of more than 100,000 (including putatively paralogous contigs) or 10,000 (excluding putatively paralogous contigs) SNPs in around 10,000 and 4000 (mostly) annotated contigs/genes, respectively. These numbers are similar to a RAD-Seq or genotyping-by-sequencing (GBS) approach (Davey *et al.* 2011), with the advantage of targeting functional, often non-anonymous regions in the genome with few missing data. Exome capture, however, is certainly more costly than RAD-Seq or GBS (but see Puritz & Lotterhos in press). In our case, this is counterbalanced by the use of DNA population pools. It is important to note that, similar to candidate gene sequencing, the genotyped set of SNPs might not be representative for the genome and is therefore biased towards, e.g., a lower proportion of neutral sites compared to whole-genome re-sequencing or alternative genome-wide genotyping.

Our data also clearly show that the probes designed for a given study species can be applied to a closely related species. For *P. cembra* and *P. sibirica*, almost the same number of contigs had mapped reads with enough coverage for reliable SNP calling, and most of these contigs were shared among both species (Table 2). The divergence time of these taxa is unclear and reported within a range of 10,000 (Krutovsky *et al.* 1994) to several million years (Saladin *et al.* 2017). However, our results show that the divergence of the two species has not reached a level of genetic differences that would have led to selectivity of the probes. Similar results have been shown in various species (for examples see Jones & Good 2016) including conifers (Neves *et al.* 2013; Suren *et al.* 2016). This opens possibilities for future investigations, such as the comparison of neutral and adaptive genetic variation in both species and their role in a recent or ongoing speciation event. The *P. sibirica* pools showed much higher genetic diversity than those of *P. cembra*, most likely because *P. cembra* pools represent discrete populations from a small area, whereas *P. sibirica* samples are consisting of bulked seeds coming from a large geographic region.

### Identification of paralogous contigs

Analyses of the two populations with individual data showed a clear heterozygote excess and a skewed AFS (Fig. 2). We interpret this as a massive presence of paralogous genes, hence probes hybridized to multiple sites in the genome that slightly differ in nucleotide sequence, but are mapped to the same contig in the bioinformatic process. It is well known that pine genomes are highly repetitive, mostly due to a high frequency of retrotransposons (Morse *et al.* 2009) and to a lesser extent due to an early WGD event (Li *et al.* 2015). Without a well-assembled reference genome, it is nearly impossible to distinguish between the different copies, making it necessary to a posteriori filter out SNPs and contigs with signs of paralogous state. Unfortunately, this requires to exclude large parts of the genome, not only of single genes, but also of certain regions such as the distal regions of the chromosomes, which are known to have a higher proportion of multi-copy genes and an important role in adaptation (Limborg *et al.* 2016).

In our case, we removed 54 and 85% of the contigs and SNPs, respectively, when looking at the two thoroughly investigated populations (EN-HJ and WC-HJ), and 63 and 87%, respectively, when considering all seven populations. Most contigs displayed the same pattern in both EN-HJ and WC-HJ (Table S9), showing that the conservatively defined set of 4950 single-copy contigs can reliably be used in other populations. The proportion of removed SNPs is very similar to that reported by Yeaman *et al.* (2016) in two North-American conifer species, for which the SNP set (Suren *et al.* 2016) was reduced by an order of magnitude. The discrepancy in the proportion of removed SNPs versus contigs comes from the fact that paralogous contigs generally contain more SNPs, and the probability of detecting multi-copy contigs increases with the number of SNPs. It is very important to note that without individual genotype data (i.e. only Pool-Seq data), we would not have been able to recognize and deal with the problem of multi-copy loci.

### Effect of paralogous contigs on patterns of neutral and adaptive genetic variation

Comparing the full (including single- and multi-copy) and single-copy SNP datasets for characterizing neutral and adaptive genetic variation revealed that ignoring the existence of paralogous contigs can have substantial consequences on the outcomes and interpretation of analyses. At first sight, the neutral genetic structure was similar using either the full or single-copy dataset (Fig. 4). The Western and Eastern genetic clusters of Swiss *P. cembra* (Gugerli *et al.,* unpublished data) could be clearly distinguished with both datasets. However, results were not that congruent for further substructure. Using simulated datasets, Willis *et al.* (2017) showed that ignoring paralogous loci can have dramatic consequences on the outcome of population structure analysis. In the present study, population genetic parameters like *H*_e_ and *F*_ST_ basically correlated well between both datasets. However, ignoring the presence of paralogous contigs led to substantial overestimation of within-population diversity (*H*_e_) and consequently underestimated among-population differentiation (*F*_ST_) due to the high level of heterozygosity of SNPs, individuals, and populations. Therefore, SNP datasets that are not filtered for paralogous loci should be treated with caution for assessing genetic diversity and differentiation. This is the case for any sequencing/genotyping approach that is not based on a high-quality reference genome.

To our knowledge, our study is the first to test the effect of removing putative paralogous contigs on investigations of adaptive genetic variation. Using an EAA (LFMM), we show that 10-30 times fewer significant associations can be found in the single-copy dataset compared to the full dataset. Since the full dataset is about 10 times larger than the single-copy dataset, most of this difference can be explained by the sheer number of possible SNPs to be associated environmental factors. On the other hand, the large number of associations is surprising, because one could expect that SNPs with a high degree of heterozygosity (and therefore mostly intermediate allele frequencies) would have a smaller chance to be detected by EAA than SNPs that have large allele frequency differences in populations from environmentally contrasting habitats. In most cases, all significant associations of the single-copy dataset were also found in the full dataset, and the *z* scores of the shared SNPs were highly correlated (Fig. S4). This robustness of LFMM to the different datasets comes from the fact that neutral genetic structure, which is incorporated as a random factor, was very similar for both datasets (see above). Nevertheless, this does not change the fact that using multi-copy SNPs in EAA will lead to spurious associations that are based on wrong allele frequencies.

### Conclusions and future directions

Our study shows how the combination of transcriptome sequencing and exome capture can be used to study neutral and adaptive genetic variation in non-model species with large genomes, but also emphasizes that the failure of avoiding or removing multi-copy loci will likely lead to erroneous results and conclusions in population genetic studies. This is the case for any sequencing/genotyping approach that is not based on a high-quality reference genome. Therefore, we consider establishing a high-quality reference genome to be essential, because it facilitates, for example, the design of probes, the identification of paralogous regions, and structural and functional annotation. However, beside the sheer size of the genome, which represents economical and computational challenges, it is the repetitive nature itself of such large genomes that has hindered and still prevents the establishment of high-quality reference genomes (Neale & Kremer 2011). To identify multi-copy loci, DNA of haploid tissue, such as the maternal megagametophyte in conifers, could be used in a sequencing or genotyping approach, because only multi-copy loci may exhibit a heterozygote state (Limborg *et al.* 2016). Methodological improvements towards this direction will help avoiding that species with large and complex genomes are excluded from studies of local adaptation using genomic approaches.

## Acknowledgements

We thank Lolita Ammann, Diego Galvan, René Graf, Lena Hellmann, Stefan Klesse, Janina Müller, Max Schmid, and Robin Winiger for support in the field; Konstantin Krutovsky and Elena A. Shilkina for providing *P. sibirica* seeds; Simon Dummermuth, Kevin Kleeb, Claudia Michel, and Christoph Sperisen for help or advice in the laboratory; Dirk Schmatz for support in extracting climate data; Olivier François, Garrett McKinney, Jason Holliday, Anne-Marie Waldvogel, and Sam Yeaman for ideas and discussion; and forest owners for sampling permission. This study was funded by the Swiss National Science Foundation (31003A_152664/1) awarded to F.G.

## Data accessibility

Raw reads used for the transcriptome assembly and SNP calling will be made available at the European Nucleotide Archive (ENA) upon acceptance. The transcriptome assembly will be deposited at the NCBI GenBank upon acceptance. Other raw data (allele frequencies and probe sequences) will be uploaded to Dryad upon acceptance.

## Author contributions

F.G. acquired funding. C.R., F.G., S.B., and S.Z. designed the study. S.B. and C.R. planned and performed the sampling. S.B. performed the laboratory work. S.Z. performed the bioinformatic analyses. B.D. prepared the topo-climatic data. C.R. und B.D. analyzed the data. C.R. wrote the manuscript, with major contributions from B.D. and F.G. All authors read, commented, and approved the current version of the article.

## References

Baker EAG, Wegrzyn JL, Sezen UU, et al. (2018) Comparative transcriptomics among four white pine species. G3: Genes|Genomes|Genetics, 8, 1461.

Bamshad MJ, Ng SB, Bigham AW, et al. (2011) Exome sequencing as a tool for Mendelian disease gene discovery. Nature Reviews Genetics, 12, 745–755.

Bansal V, Tewhey R, LeProust EM, Schork NJ (2011) Efficient and cost effective population resequencing by pooling and in-solution hybridization. PLOS ONE, 6, e18353.

Barth JMI, Berg PR, Jonsson PR, et al. (2017) Genome architecture enables local adaptation of Atlantic cod despite high connectivity. Molecular Ecology, 26, 4452–4466.

Benjamini Y, Hochberg Y (1995) Controlling the false discovery rate – a practical and powerful approach to multiple testing. Journal of the Royal Statistical Society Series B-Methodological, 57, 289–300.

Bolger AM, Lohse M, Usadel B (2014) Trimmomatic: a flexible trimmer for Illumina sequence data. Bioinformatics, 30, 2114–2120.

Caudullo G, Welk E, San-Miguel-Ayanz J (2017) Chorological maps for the main European woody species. Data in Brief, 12, 662–666.

Caye K, François O (2018) LFMM 2.0: Latent factor models for confounder adjustment in genome and epigenome-wide association studies. bioRxiv, 255893.

Chen H, Boutros PC (2011) VennDiagram: a package for the generation of highly-customizable Venn and Euler diagrams in R. BMC Bioinformatics, 12, 7.

Clark LV, Jasieniuk M (2011) POLYSAT: an R package for polyploid microsatellite analysis. Molecular Ecology Resources, 11, 562–566.

Csillery K, Rodriguez-Verdugo A, Rellstab C, Guillaume F (2018) Detecting the genomic signal of polygenic adaptation and the role of epistasis in evolution. Molecular Ecology, 27, 606–612.

Davey JW, Hohenlohe PA, Etter PD, et al. (2011) Genome-wide genetic marker discovery and genotyping using next-generation sequencing. Nature Reviews Genetics, 12, 499–510.

de Villemereuil P, Frichot E, Bazin E, François O, Gaggiotti OE (2014) Genome scan methods against more complex models: when and how much should we trust them. Molecular Ecology, 23, 2006–2019.

De Wit P, Palumbi SR (2013) Transcriptome-wide polymorphisms of red abalone (Haliotis rufescens) reveal patterns of gene flow and local adaptation. Molecular Ecology, 22, 2884–2897.

Eckert AJ, Bower AD, González-Martínez SC, et al. (2010) Back to nature: ecological genomics of loblolly pine (Pinus taeda, Pinaceae). Molecular Ecology, 19, 3789–3805.

Exposito-Alonso M, Vasseur F, Ding W, et al. (2018) Genomic basis and evolutionary potential for extreme drought adaptation in Arabidopsis thaliana. Nature Ecology & Evolution, 2, 352–358.

Fischer MC, Rellstab C, Leuzinger M, et al. (2017) Estimating genomic diversity and population differentiation – an empirical comparison of microsatellite and SNP variation in Arabidopsis halleri. BMC Genomics, 18, 69.

Fischer MC, Rellstab C, Tedder A, et al. (2013) Population genomic footprints of selection and associations with climate in natural populations of Arabidopsis halleri from the Alps. Molecular Ecology, 22, 5594–5607.

Flood PJ, Hancock AM (2017) The genomic basis of adaptation in plants. Current Opinion in Plant Biology, 36, 88–94.

François O, Martins H, Caye K, Schoville SD (2016) Controlling false discoveries in genome scans for selection. Molecular Ecology, 25, 454–469.

Frichot E, Schoville SD, Bouchard G, François O (2013) Testing for associations between loci and environmental gradients using latent factor mixed models. Molecular Biology and Evolution, 30, 1687–1699.

Gernandt DS, Lopez GG, Garcia SO, Liston A (2005) Phylogeny and classification of Pinus. Taxon, 54, 29–42.

Götz S, García-Gomez JM, Terol J, et al. (2008) High-throughput functional annotation and data mining with the Blast2GO suite. Nucleic Acids Research, 36, 3420–3435.

Grabherr MG, Haas BJ, Yassour M, et al. (2011) Full-length transcriptome assembly from RNA-Seq data without a reference genome. Nature Biotechnology, 29, 644–U130.

Günther T, Coop G (2013) Robust identification of local adaptation from allele frequencies. Genetics, 195, 205–220.

Günther T, Lampei C, Barilar I, Schmid KJ (2016) Genomic and phenotypic differentiation of Arabidopsis thaliana along altitudinal gradients in the North Italian Alps. Molecular Ecology, 25, 3574–3592.

Hoban S, Kelley JL, Lotterhos KE, et al. (2016) Finding the genomic basis of local adaptation: pitfalls, practical solutions, and future directions. American Naturalist, 188, 379–397.

Hohenlohe PA, Phillips PC, Cresko WA (2010) Using population genomics to detect selection in natural populations: key concepts and methodological considerations. International Journal of Plant Sciences, 171, 1059–1071.

Jones MR, Good JM (2016) Targeted capture in evolutionary and ecological genomics. Molecular Ecology, 25, 185–202.

Jones P, Binns D, Chang HY, et al. (2014) InterProScan 5: genome-scale protein function classification. Bioinformatics, 30, 1236–1240.

Korte A, Farlow A (2013) The advantages and limitations of trait analysis with GWAS: a review. Plant Methods, 9, 29.

Krutovsky KV, Politov DV, Altukhov YP (1994) Genetic differentiation and phylogeny of stone pine species based on isozyme loci. In: Proceedings - International workshop on subalpine stone pints and their environment: the status of our knowledge (eds. Schmidt WC, Holtmeier F-K), pp. 19–30.

Langmead B, Trapnell C, Pop M, Salzberg SL (2009) Ultrafast and memory-efficient alignment of short DNA sequences to the human genome. Genome Biology, 10, 10.

Laporte M, Pavey SA, Rougeux C, et al. (2016) RAD sequencing reveals within-generation polygenic selection in response to anthropogenic organic and metal contamination in North Atlantic Eels. Molecular Ecology, 25, 219–237.

Li Z, Baniaga AE, Sessa EB, et al. (2015) Early genome duplications in conifers and other seed plants. Science Advances, 1, e1501084.

Limborg MT, Seeb LW, Seeb JE (2016) Sorting duplicated loci disentangles complexities of polyploid genomes masked by genotyping by sequencing. Molecular Ecology, 25, 2117–2129.

Lotterhos KE, Whitlock MC (2015) The relative power of genome scans to detect local adaptation depends on sampling design and statistical method. Molecular Ecology, 24, 1031–1046.

Machado HE, Bergland AO, O’Brien KR, et al. (2016) Comparative population genomics of latitudinal variation in Drosophila simulans and Drosophila melanogaster. Molecular Ecology, 25, 723–740.

Mantel N (1967) Detection of disease clustering and a generalized regression approach. Cancer Research, 27, 209–220.

McKenna A, Hanna M, Banks E, et al. (2010) The Genome Analysis Toolkit: A MapReduce framework for analyzing next-generation DNA sequencing data. Genome Research, 20, 1297–1303.

McKinney GJ, Waples RK, Seeb LW, Seeb JE (2016) Data from: Paralogs are revealed by proportion of heterozygotes and deviations in read ratios in genotyping by sequencing data from natural populations. Dryad Data Repository. https://doi.org/10.5061/dryad.cm08m

McKinney GJ, Waples RK, Seeb LW, Seeb JE (2017) Paralogs are revealed by proportion of heterozygotes and deviations in read ratios in genotyping-by-sequencing data from natural populations. Molecular Ecology Resources, 17, 656–669.

Meier ES, Kienast F, Pearman PB, et al. (2010) Biotic and abiotic variables show little redundancy in explaining tree species distributions. Ecography, 33, 1038–1048.

Morgulis A, Gertz EM, Schaffer AA, Agarwala R (2006) A fast and symmetric DUST implementation to mask low-complexity DNA sequences. Journal of Computational Biology, 13, 1028–1040.

Morse AM, Peterson DG, Islam-Faridi MN, et al. (2009) Evolution of genome size and complexity in Pinus. PLOS ONE, 4, e4332.

Mosca E, Gugerli F, Eckert AJ, Neale DB (2016) Signatures of natural selection on Pinus cembra and P. mugo along elevational gradients in the Alps. Tree Genetics & Genomes, 12, 9.

Murray B, Leitch I, Bennett M (2012) Gymnosperm DNA C-values Database (Release 5.0, Dec. 2012). http://data.kew.org/cvalues/

Neale DB, Kremer A (2011) Forest tree genomics: growing resources and applications. Nature Reviews Genetics, 12, 111–122.

Neuschulz EL, Merges D, Bollmann K, Gugerli F, Böhning-Gaese K (2018) Biotic interactions and seed deposition rather than abiotic factors determine recruitment at elevational range limits of an alpine tree. Journal of Ecology, 106, 948–959.

Neves LG, Davis JM, Barbazuk WB, Kirst M (2013) Whole-exome targeted sequencing of the uncharacterized pine genome. Plant Journal, 75, 146–156.

Oksanen J, Blanchet FG, Kindt R, et al. (2013) vegan: Community Ecology Package. R package version 2.0-8. http://CRAN.R-project.org/package=vegan

Pinosio S, González-Martínez SC, Bagnoli F, et al. (2014) First insights into the transcriptome and development of new genomic tools of a widespread circum-Mediterranean tree species, Pinus halepensis Mill. Molecular Ecology Resources, 14, 846–856.

Pritchard JK, Di Rienzo A (2010) Adaptation — not by sweeps alone. Nature Reviews Genetics, 11, 665–667.

Puritz JB, Lotterhos KE (in press) Expressed exome capture sequencing: A method for cost-effective exome sequencing for all organisms. Molecular Ecology Resources, doi: 10.1111/1755-0998.12905.

Quinlan AR, Hall IM (2010) BEDTools: A flexible suite of utilities for comparing genomic features. Bioinformatics, 26, 841–842.

R Development Core Team (2018) R: a language and environment for statistical computing. http://www.R-project.org

Rellstab C, Gugerli F, Eckert A, Hancock A, Holderegger R (2015) A practical guide to environmental association analysis in landscape genomics. Molecular Ecology, 24, 4348–4370.

Rellstab C, Zoller S, Tedder A, Gugerli F, Fischer MC (2013) Validation of SNP allele frequencies determined by pooled next-generation sequencing in natural populations of a non-model plant species. PLOS ONE, 8, e80422.

Rellstab C, Zoller S, Walthert L, et al. (2016) Signatures of local adaptation in candidate genes of oaks (Quercus spp.) with respect to present and future climatic conditions. Molecular Ecology, 25, 5907–5924.

Ryu S, Han J, Norden-Krichmar TM, Schork NJ, Suh Y (2018) Effective discovery of rare variants by pooled target capture sequencing: a comparative analysis with individually indexed target capture sequencing. Mutation Research/Fundamental and Molecular Mechanisms of Mutagenesis, 809, 24–31.

Saladin B, Leslie AB, Wuest RO, et al. (2017) Fossils matter: improved estimates of divergence times in Pinus reveal older diversification. BMC Evolutionary Biology, 17, 95.

Salzer K (2011) Wind-and bird-mediated gene flow in Pinus cembra: Effects on spatial genetic structure and potential close-relative inbreeding (PhD Thesis), University of Zürich.

Salzer K, Gugerli F (2012) Reduced fitness at early life stages in peripheral versus core populations of Swiss stone pine (Pinus cembra) is not reflected by levels of inbreeding in seed families. Alpine Botany, 122, 75–85.

Schlötterer C, Tobler R, Kofler R, Nolte V (2014) Sequencing pools of individuals – mining genome-wide polymorphism data without big funding. Nature Review Genetics, 15, 749–763.

Stevens KA, Wegrzyn JL, Zimin A, et al. (2016) Sequence of the sugar pine megagenome. Genetics, 204, 1613–1626.

Suren H, Hodgins KA, Yeaman S, et al. (2016) Exome capture from the spruce and pine giga-genomes. Molecular Ecology Resources, 16, 1136–1146.

Tiffin P, Ross-Ibarra J (2014) Advances and limits of using population genetics to understand local adaptation. Trends in Ecology & Evolution, 29, 673–680.

Wachowiak W, Trivedi U, Perry A, Cavers S (2015) Comparative transcriptomics of a complex of four European pine species. BMC Genomics, 16, 9.

Waldvogel A-M, Wieser A, Schell T, et al. (2017) The genomic footprint of climate adaptation in Chironomus riparius. Molecular Ecology, 27, 1439–1456.

Wegrzyn JL, Liechty JD, Stevens KA, et al. (2014) Unique features of the loblolly pine (*Pinus taeda L*.) megagenome revealed through sequence annotation. Genetics, 196, 891–909.

Weir BS, Cockerham CC (1984) Estimating F-statistics for the analysis of population structure. Evolution, 38, 1358–1370.

Willis SC, Hollenbeck CM, Puritz JB, Gold JR, Portnoy DS (2017) Haplotyping RAD loci: an efficient method to filter paralogs and account for physical linkage. Molecular Ecology Resources, 17, 955–965.

Yamada K, Kanai M, Osakabe Y, et al. (2011) Monosaccharide absorption activity of Arabidopsis roots depends on expression profiles of transporter genes under high salinity conditions. Journal of Biological Chemistry, 286, 43577–43586.

Yeaman S, Hodgins KA, Lotterhos KE, et al. (2016) Convergent local adaptation to climate in distantly related conifers. Science, 353, 1431–1433.

Zimin A, Stevens KA, Crepeau M, et al. (2014) Sequencing and assembly of the 22-Gb loblolly pine genome. Genetics, 196, 875–890.

Zonneveld BJM (2012) Conifer genome sizes of 172 species, covering 64 of 67 genera, range from 8 to 72 picogram. Nordic Journal of Botany, 30, 490–502.

